# High temperature treatment optimized for symbiont suppression in an obligatory gut bacterial symbiosis in the stinkbug *Plautia stali*

**DOI:** 10.1101/2024.04.04.588189

**Authors:** Wen-Jin Cai, Minoru Moriyama, Takema Fukatsu

**Author notes:** Correspondence: Minoru Moriyama, Takema Fukatsu.

## Abstract

In the era of global warming, much attention has been paid to the possibility that high temperature may influence diverse insects not only directly but also indirectly via effects on their symbiotic microorganisms. The stinkbug *Plautia stali* develops a midgut symbiotic organ that harbors a specific bacterial symbiont indispensable for its growth and survival. Being maintainable in laboratory and tractable experimentally, *P. stali* is recently highlighted as a model system to investigate the mechanisms underpinning insect-microbe symbiotic interactions. In this study, we reared newly-emerged adult insects of *P. stali* under different temperature conditions for 8 days and monitored how their symbiotic organs and symbiotic bacteria are affected. While all insects survived at temperatures from 25°C to 37°C, some insects died at 38°C, 39°C and 40°C, wherein mortality rates increased as temperature elevated. While the symbiotic organs of the normal insects exhibited vivid yellow color, the symbiotic organs of the insects reared at 35°C or higher frequently exhibited abnormal colors such as pale yellow, yellowish white, or white, the extent of which became more severe as temperature elevated. Symbiont quantification revealed that, while the symbiont titers were almost constant for 8 days at 25°C and 30°C, the symbiont titers on the 8^th^ day drastically declined to 1/100 at 35℃ and 1/10000 at 37°C and 39°C. Based on these results, we propose that rearing at 37°C for a week is a recommended treatment regime by which the symbiont is effectively suppressed with minimal damage to the host insect.

## Introduction

In the era of global warming, it has become a worldwide concern that elevated temperature associated with global climate change deteriorates the ecosystem and the biodiversity (Walther 2010; Gilman et al. 2010; Harley 2011), where effects by way of inter-organismal interactions may play significant roles (Kiers et al. 2010; Blois et al. 2013; Altizer et al. 2013). Considering that many insects are intimately associated with symbiotic microorganisms (Buchner 1965; Bourtzis and Miller 2003; Douglas 2022), it is important to understand how such microbial associates are involved in deterioration, tolerance and/or adaptation of insects to high temperature conditions (Renoz et al. 2019; Wernegreen 2012; Rodrigues & Beldade 2020; Lemoine et al. 2020).

Previous studies have uncovered that high temperature treatments tend to suppress or even eliminate microbial symbionts, and such aposymbiotic insects often suffer negative fitness consequences as reported in bedbugs (Chang 1974), planthoppers (Noda and Saito 1979), aphids (Ohtaka and Ishikawa 1991), cockroaches (Sacchi et al. 1993), stinkbugs (Prado et al. 2010), ants (Fan and Wernegreen 2013), whiteflies (Shan et al. 2017), weevils (Anbutsu et al. 2017) and others. Simulated global warming with reference to natural ambient temperatures resulted in suppression of obligatory gut symbiotic bacteria, abnormal body coloration, and remarkable growth retardation in the stinkbug *Nezara viridula* (Kikuchi et al. 2016). In the pea aphid *Acyrthosiphon pisum*, some facultative bacterial symbionts like *Serratia symbiotica* confer host’s resistance to heat stress (Montllor et al. 2002; Russell and Moran 2006), and some genotypes of the obligatory bacterial symbiont *Buchnera aphidicola* affect host’s tolerance to high temperature conditions (Dunbar et al. 2007; Zhang et al. 2019). As such, symbiotic bacteria are deeply involved in physiological, ecological and evolutionary responses of host insects to high temperature conditions, but what molecular mechanisms underlie the processes has been poorly understood, mainly because both the host and the symbiont are genetically intractable in most of the insect-microbe symbiotic systems.

In this context, the brown-winged green stinkbug *Plautia stali* Scott (Hemiptera: Pentatomidae) (Fig. 1a) has recently emerged as a promising model system in that (i) the insect has a well-developed midgut organ (Fig. 1b) to harbor specific gut symbiotic bacteria of the genus *Pantoea* essential for growth and survival (Abe et al. 1995; Hosokawa et al. 2016), (ii) the insect is easily and stably maintainable in laboratory (Nishide et al. 2017), (iii) experimental manipulations of the symbiotic association, such as symbiont removal, re-inoculation, replacement, etc., are conveniently executable (Hosokawa et al. 2016), (iv) some symbiont strains are cultivable and thus genetically manipulatable (Hosokawa et al. 2016), (v) RNA interference knockdown of host gene expression works efficiently (Futahashi et al. 2011; Nishide et al. 2020; Nishide et al. 2021; Moriyama et al. 2022), (vi) the outstanding genetic model bacterium, *Escherichia coli*, can infect and evolve into a mutualistic symbiont of the insect using experimental evolutionary approaches (Koga et al. 2022). In this study, we maintained newly-emerged adult insects of *P. stali* under different high temperature treatments in a systematic manner and evaluated the effects on the host symbiotic organ as well as the symbiotic bacteria, thereby establishing an optimal treatment scheme for effectively disrupting the microbial symbiosis with minimal damage to the host insect.

**Fig. 1.**
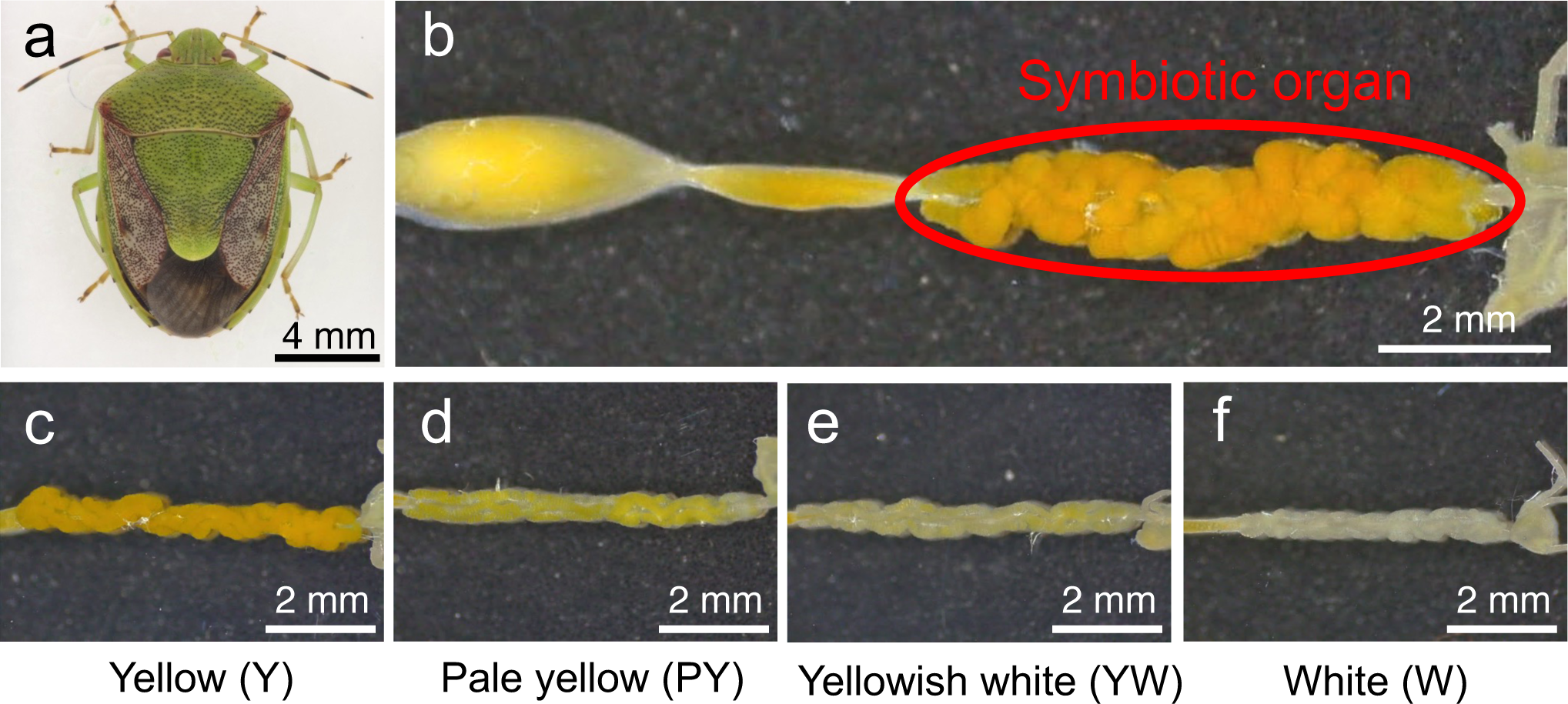
Color categories of the midgut symbiotic organ of *P. stali*. (**a**) An adult insect. (**b**) A dissected alimentary tract, in which red circle highlights the symbiotic organ. (**c**) Yellow (Y). (**d**) Pale yellow (PY). (**e**) Yellowish white (YW). (**f**) White (W).

## Materials and Methods

### Insect material

A long-lasting laboratory strain of *P. stali*, which was originally derived from adult insects collected at Tsukuba, Ibaraki, Japan in 2012, was used. The insects were maintained in plastic containers and fed with raw peanuts, raw soybeans, and distilled water supplemented with 0.05% (w/v) ascorbic acid (DWA) in climate chambers at 25 ± 1°C under a long-day regime of 16 h light and 8 h dark as described (Nishide et al. 2017).

### Temperature treatment

From the insect stock maintained as described above, sixteen adults 1 day after eclosion were collected and transferred to each new rearing container. In total, 128 insects consisting of 64 males in 4 containers and 64 females in 4 containers were subjected to each temperature treatment. Setting of rearing incubators (TOMY, CLE305 or PHCBI, MLR-352) was either at 25°C, 30°C, 35°C, 36°C, 37°C, 38°C, 39°C or 40°C under a long-day regime of 16 h light and 8 h dark. Survival of the insects in each rearing container was checked every day around 10 am when DWA bottle was renewed.

### Dissection and observation of symbiotic organ

For each temperature treatment group every day, four adult males and four adult females were individually subjected to dissection of the symbiotic organ in a Petri dish filled with a phosphate-buffered saline (PBS: 137 mM NaCl, 8.1 mM Na_2_HPO_4_, 2.7 mM KCl, 1.5 mM KH_2_PO_4_, pH 7.4) using fine scissors, forceps and razor blades. The dissected symbiotic organs were photographed under a stereo-microscope (Leica, S9i), classified into the visual color categories Y (yellow), PY (pale yellow), YW (yellowish white) or W (white) (Fig. 1c-f), and then stored in an ultracold freezer at –80°C.

### Symbiont quantification

Each of the dissected symbiotic organs was homogenized in a plastic tube with a pestle in 200 μl of PBS and subjected to an alkaline DNA extraction procedure essentially as described (Truett et al. 2000). An aliquot of the homogenate (10 μl) was transferred to a new plastic tube, combined and mixed with 32 μl of alkaline solution (25 mM NaOH, 0.2 mM EDTA), heated at 95°C for 10 min, neutralized by adding and mixing with 64 μl of neutralizing solution (12.5 mM HCl, 10 mM Tris), and placed on ice. Each of the sample solutions was centrifuged and subjected to quantitative PCR either after 100 times dilution with water or as was. The primers AgroL1013F (5′-ATC AGG GCG CAA TAT CTG GT-3′) and AgroL1103R (5′-CGC TCC TGC AGT TTC TCT TTG-3′), which target a 91 bp region of bacterial *groEL* gene of the symbiont of *P. stali*, were used. The PCR reaction mixtures, each 20 μl in total, containing 300 nM each of the primers, 2 × KAPA SYBR FAST qPCR Master Mix Universal (KAPA Biosystems), 5 μl of sample solution, and water, were subjected to quantitative PCR under a temperature profile of 95°C for 3 min followed by 40 cycles for 95°C for 5 sec and 58°C for 15 sec using the Stratagene Mx3000P (Agilent Technologies). For each of the samples, two replicate measurements were conducted. Standard curves were drawn using a linearized plasmid containing the target site, with concentrations equivalent to 5.45 x 10^2^, 10^3^, 10^4^, 10^5^, 10^6^ and 10^7^ *groEL* gene copies, respectively.

## Results and Discussion

### Insect survival

When adult insects of *P. stali* were reared under different temperature conditions, all insects survived the experimental period of 8 days for 25°C, 30°C, 35°C, 36°C and 37°C treatments, whereas some insects died at higher temperatures, and mortality rates increased as temperature elevated (Fig. 2).

**Fig. 2.**
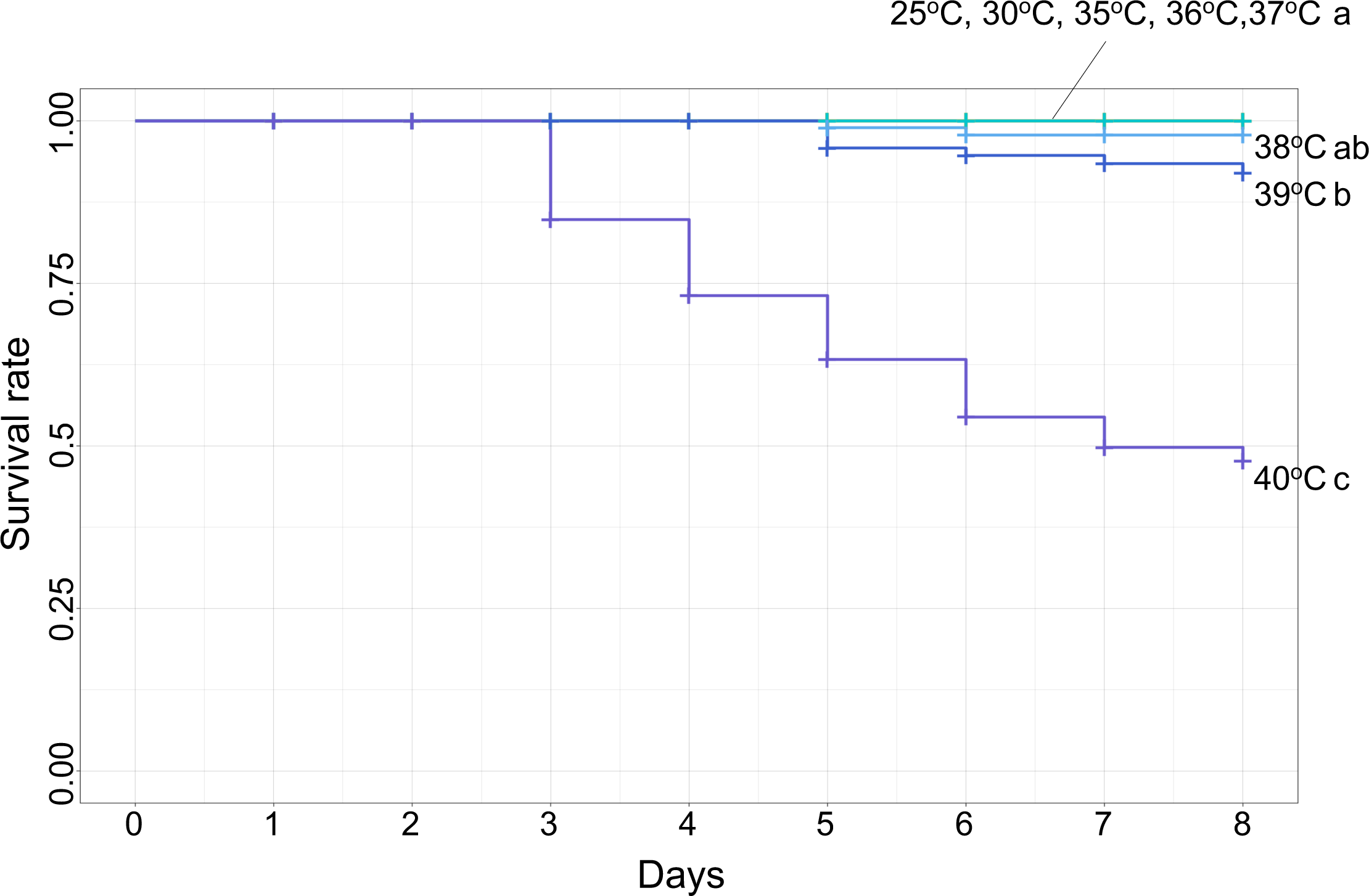
Survival curves of *P. stali* reared at different temperatures. Each treatment group started with 128 insects, of which 64 were censored for daily dissection. Different alphabetical letters (a, b, c) indicate statistically significant differences (pairwise log-rank test: *P* < 0.05 with Benjamini-hochberg correction).

### Appearance of symbiotic organ

When the insects were reared at 25°C, their symbiotic organs were constantly yellow in color, being of normal appearance throughout the experimental period of 8 days (Fig. 3a; Fig. S2). When the insects were reared at 30°C, their symbiotic organs were also constantly yellow in color, except for a case of pale yellow on the 8^th^ day (Fig. 3b; Fig. S3). When the insects were reared at 35°C, by contrast, their symbiotic organs started to exhibit abnormal appearance on the 5^th^ day and on. On the 8^th^ day, most of them were either yellowish white or white in color (Fig. 3c; Fig. S4). As the rearing temperatures were elevated to 36°C, 37°C, 38°C, 39°C and 40°C, the abnormal appearance of the symbiotic organs became more and more intense (Fig. 3d-h; Figs. S5-S9): all exhibited white or whitish yellow on the 8^th^ day at 37°C (Fig. 3e; Fig. S6), on the 7^th^ day at 38°C and 39°C (Fig. 3f, g; Figs. S7, S8), and on the 6^th^ day at 40°C (Fig. 3h; Fig. S9).

**Fig. 3.**
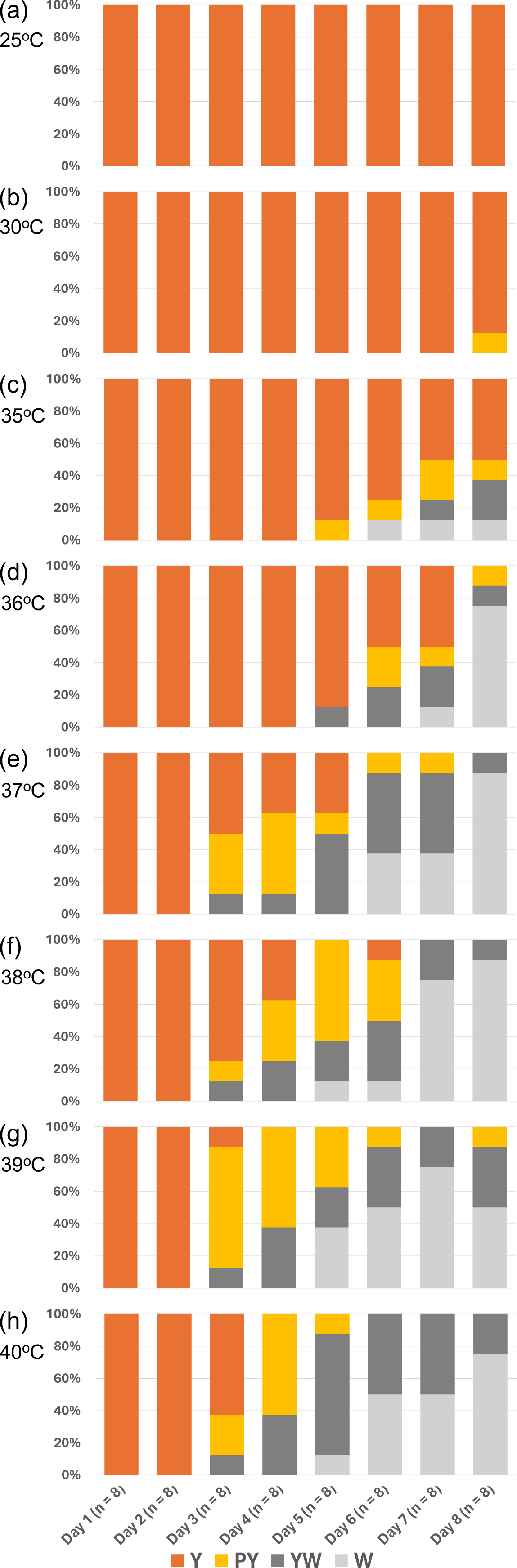
Color categories of symbiotic organs dissected from *P. stali* maintained at different high temperature conditions for 8 days. (**a**) 25°C. (**b**) 30°C. (**c**) 35°C. (**d**) 36°C. (**e**) 37°C. (**f**) 38°C. (**g**) 39°C. (**h**) 40°C.

### Titer of symbiotic bacteria

The symbiotic bacteria in the dissected symbiotic organs were quantified by quantitative PCR throughout the maintenance under the different temperature conditions. The initial symbiont titers, around 10^9^ *groEL* gene copies equivalent, were maintained throughout the experimental period of 8 days when the insects were reared at 25°C and 30°C (Fig. 4a, b). By contrast, the symbiont titers significantly declined when the host insects were reared at higher temperatures (Fig. 4c-e). On the 8^th^ day, the initial symbiont titers, around 10^9^ *groEL* gene copies equivalent, were down to 10^7^-10^8^ *groEL* gene copies equivalent at 35°C (Fig. 4c), 10^5^-10^6^ *groEL* gene copies equivalent at 37°C (Fig. 4d), and around 10^5^ *groEL* gene copies equivalent at 39°C (Fig. 4e).

**Fig. 4.**
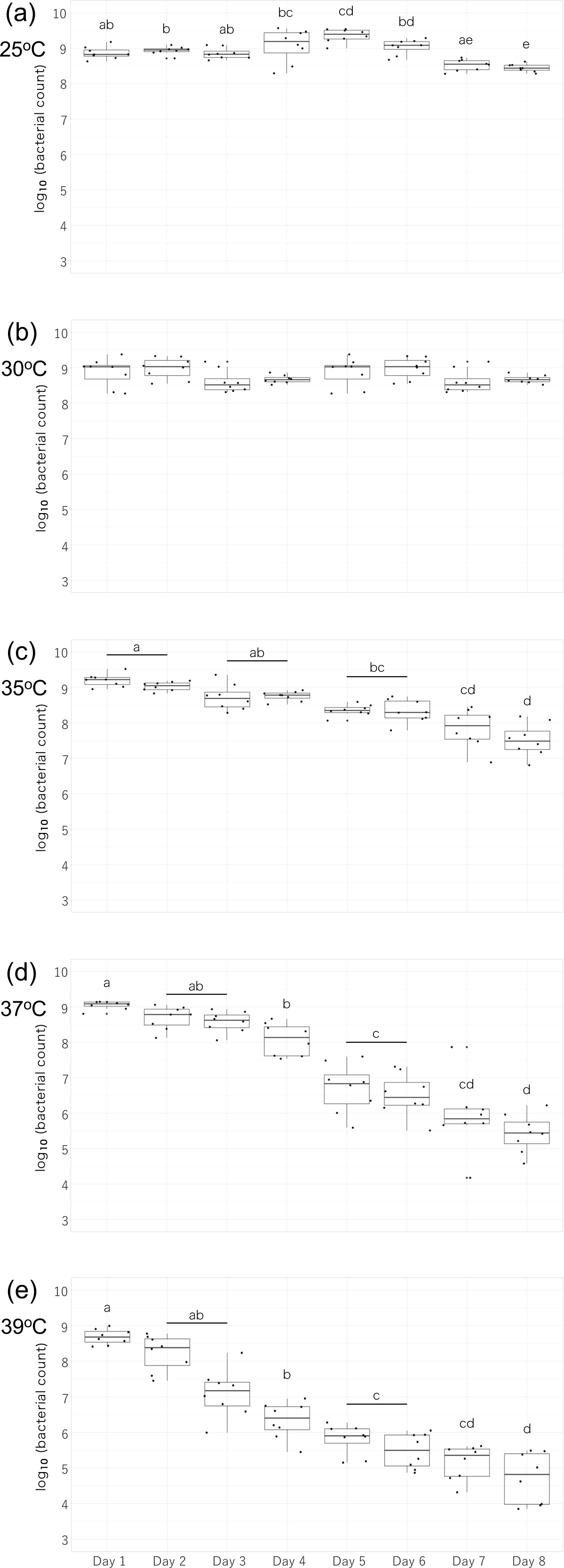
Quantitative PCR monitoring of symbiont titers during the maintenance of *P. stali* at different high temperature conditions for 8 days. (**a**) 25°C. (**b**) 30°C. (**c**) 35°C. (**d**) 37°C. (**e**) 39°C. Different alphabetical letters indicate statistically significant differences (pairwise *t*-test: *P* < 0.05 with Bonferroni correction).

### Conclusion and perspective

All these results taken together, we conclude that rearing at 37°C for a week is the best treatment scheme for experimentally disrupting the essential gut bacterial symbiosis in the stinkbug *P. stali*, by which no treated insects died, all the treated insects exhibited a colorless symbiotic organ indicative of disrupted symbiosis, and all the treated insects showed the symbiont titers around 10^5^ *groEL* gene copies, which were about 1/10000 less than those of the control insects around 10^9^ *groEL* gene copies. It seems likely that the symbiont DNA detected by quantitative PCR at very low levels represented remnant DNA derived from dead bacterial cells, although the absence of living symbiont cells is not easy to unequivocally confirm for the uncultivable bacterial symbiont.

The conspicuous disruptive effects of high ambient temperature are not restricted to insect-microbe symbiotic associations but also observed in coral-dinoflagellate mutualism (Webster and Reusch 2017), plant-pollinator mutualism (Miller-Struttmann et al. 2015), vertebrate gut microbiome (Bestion et al. 2017) and many other symbiotic associations (Kiers et al. 2010; Blois et al. 2013; Altizer et al. 2013). In this context, the gut bacterial symbiosis in the stinkbug *P. stali* provides an excellent model system that enables general understanding of the mechanisms underpinning the effects of global warming to symbiotic associations using experimental, genetic and evolutionary approaches.

## Acknowledgments

This study was supported by the Japan Science and Technology Agency (JST) ERATO grant no. JPMJER1902 to T.F.

## Supplementary Figure Captions

**Fig. S1.**
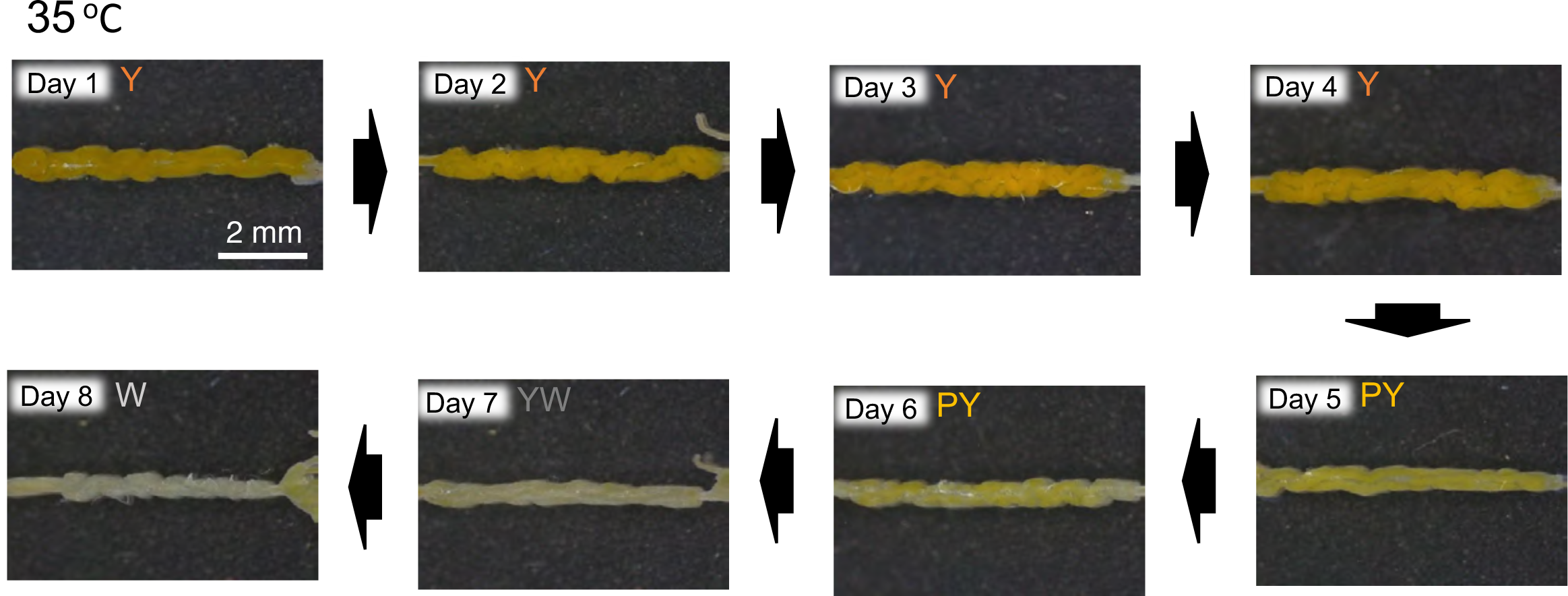
Representative images of symbiotic organs dissected from *P. stali* adults maintained at 35°C for 8 days and their color categories abbreviated as Y (yellow), PY (pale yellow), YW (yellowish white) or W (white).

**Fig S2.**
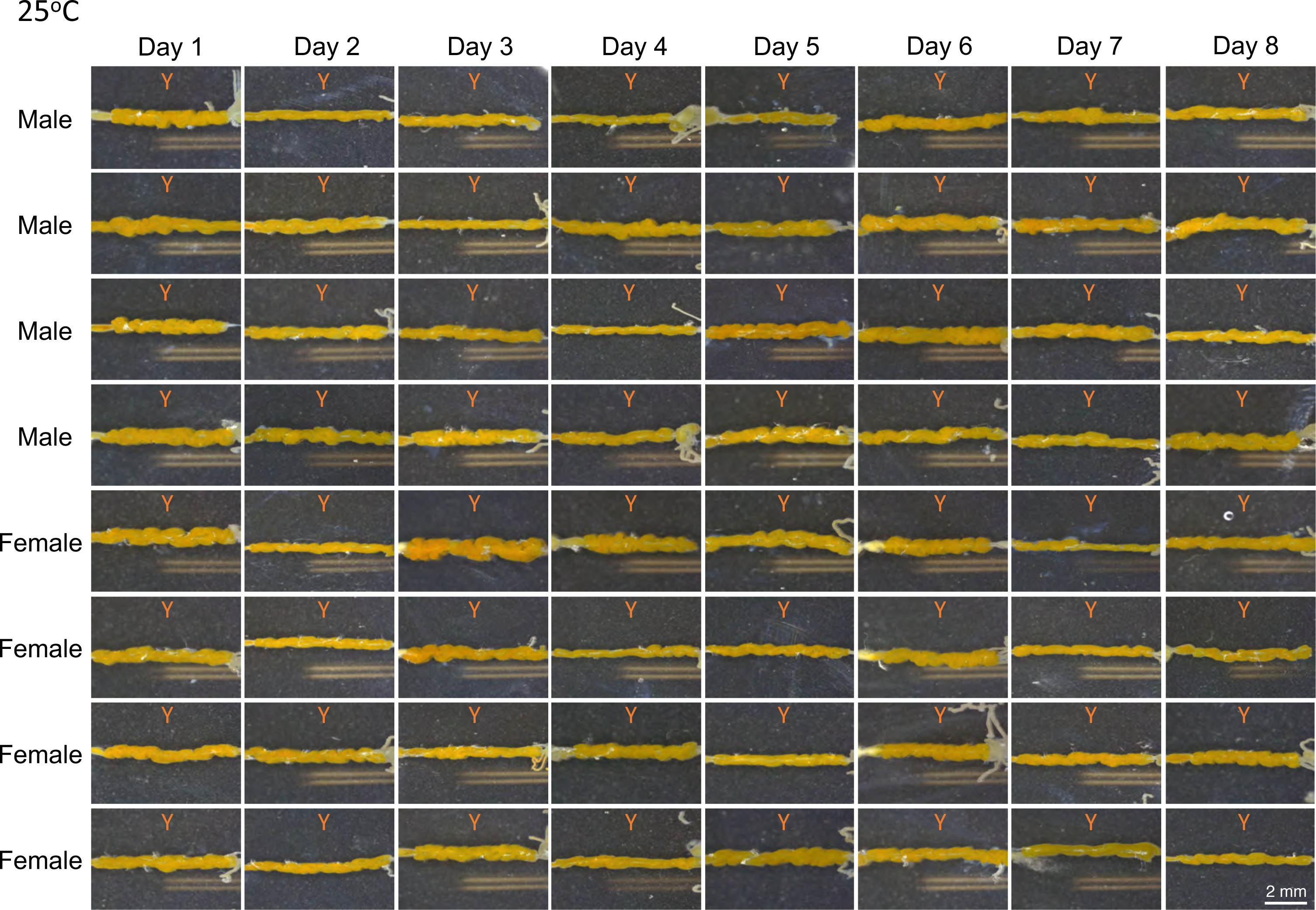
Images of symbiotic organs dissected from *P. stali* adults maintained at 25°C and their color categories abbreviated as Y (yellow). For 8 days, 4 males and 4 females were dissected and inspected every day.

**Fig S3.**
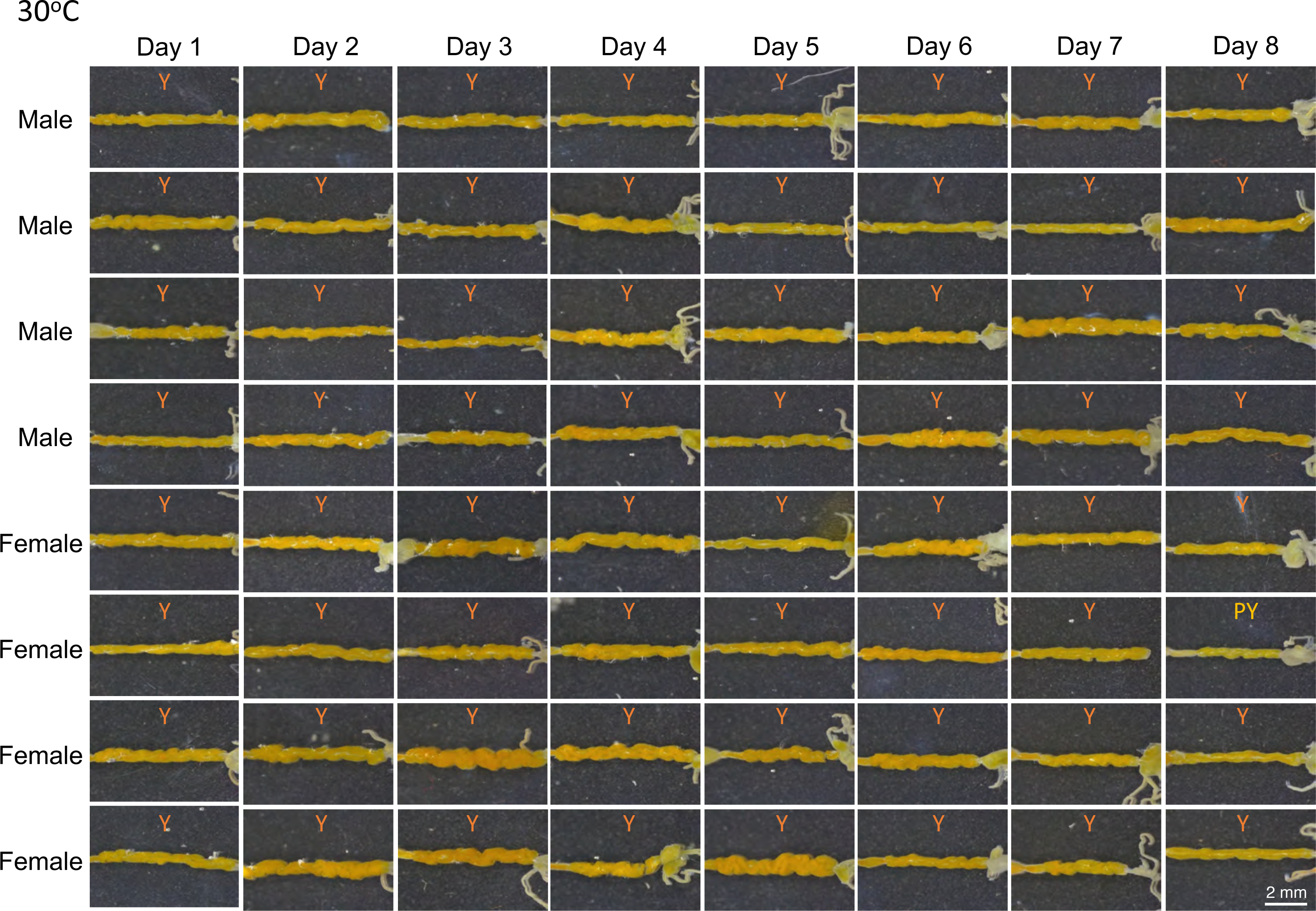
Images of symbiotic organs dissected from *P. stali* adults maintained at 30°C and their color categories abbreviated as Y (yellow) or PY (pale yellow). For 8 days, 4 males and 4 females were dissected and inspected every day.

**Fig S4.**
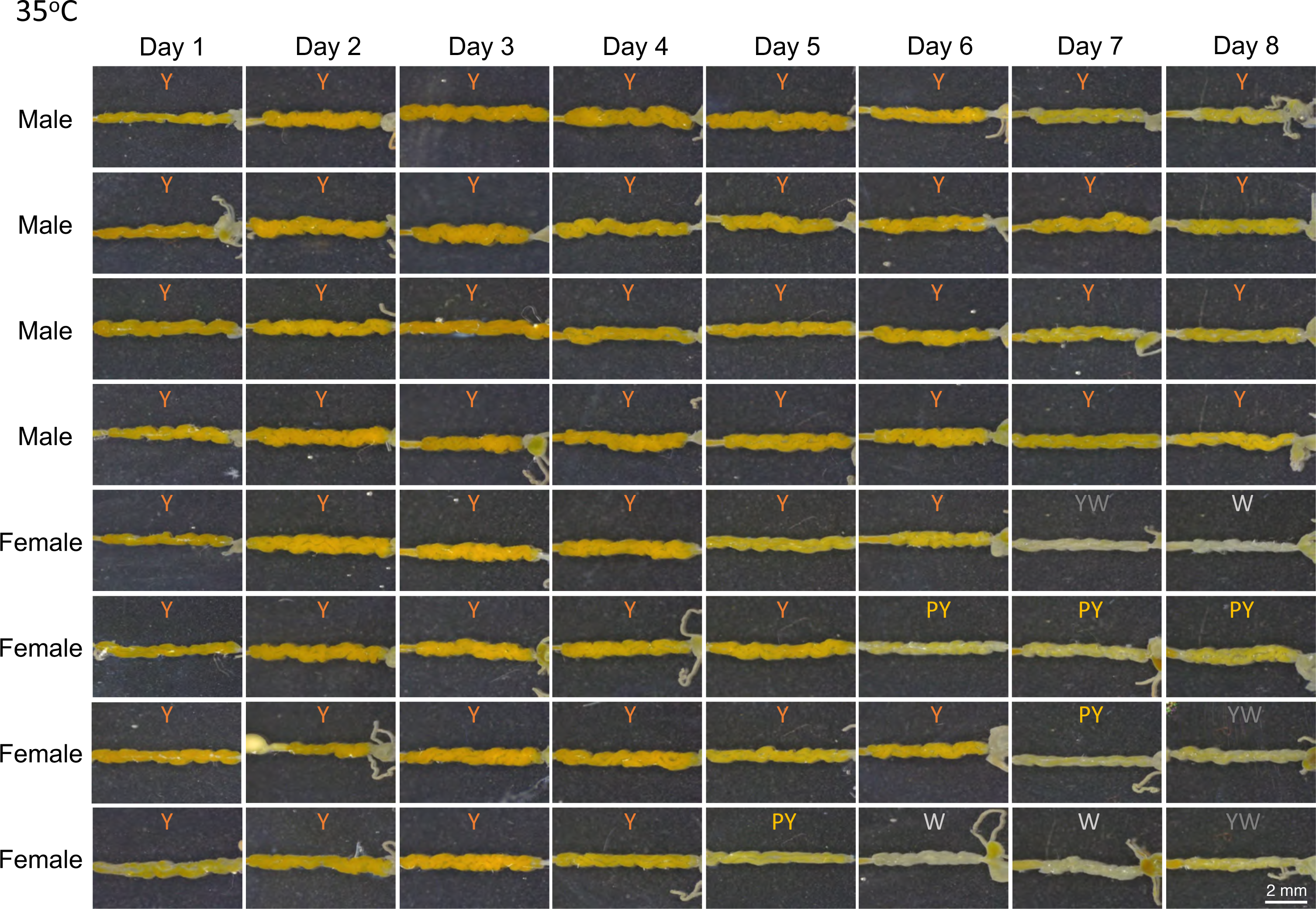
Images of symbiotic organs dissected from *P. stali* adults maintained at 35°C and their color categories abbreviated as Y (yellow), PY (pale yellow), YW (yellowish white) or W (white). For 8 days, 4 males and 4 females were dissected and inspected every day.

**Fig S5.**
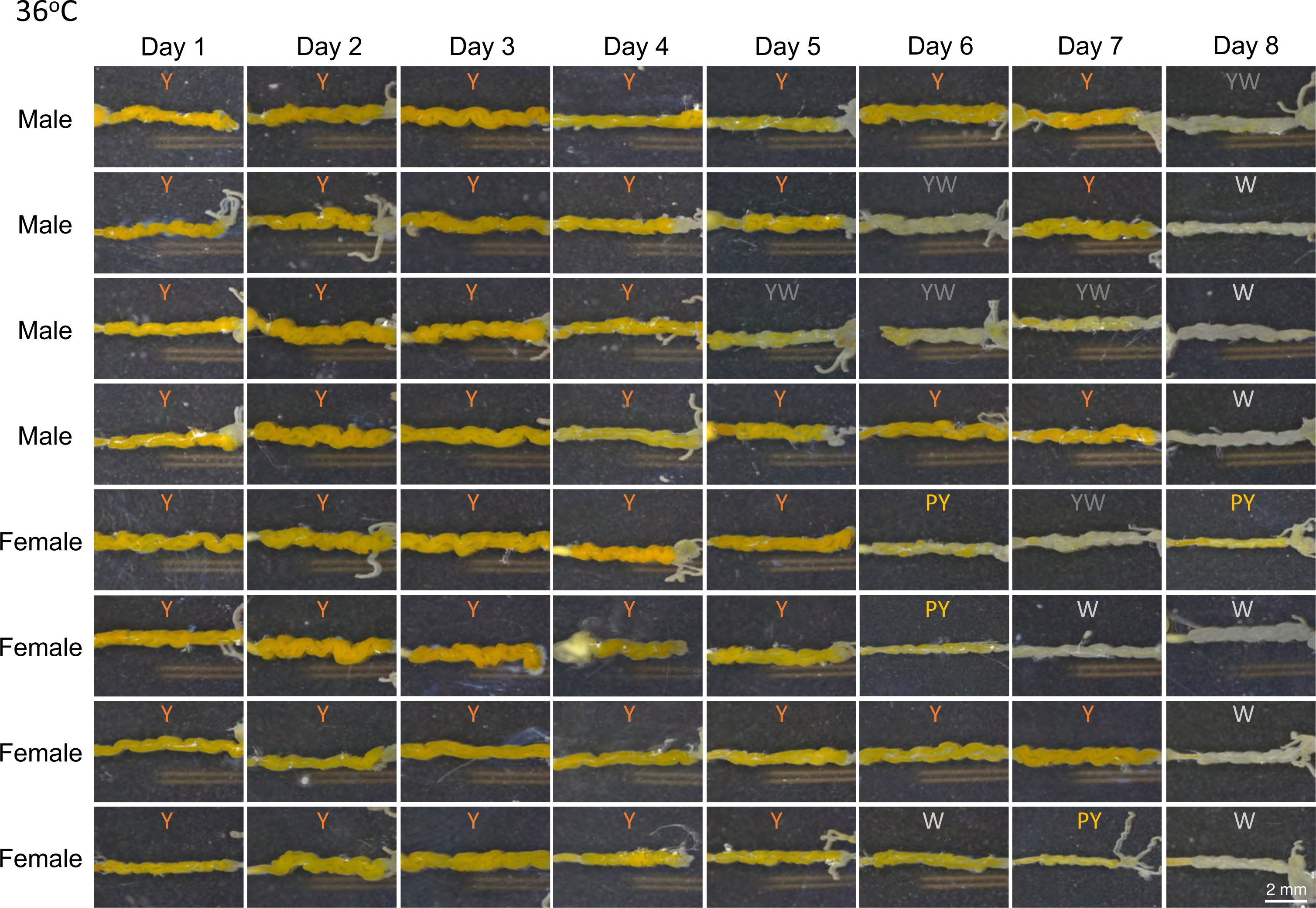
Images of symbiotic organs dissected from *P. stali* adults maintained at 36°C and their color categories abbreviated as Y (yellow), PY (pale yellow), YW (yellowish white) or W (white). For 8 days, 4 males and 4 females were dissected and inspected every day.

**Fig S6.**
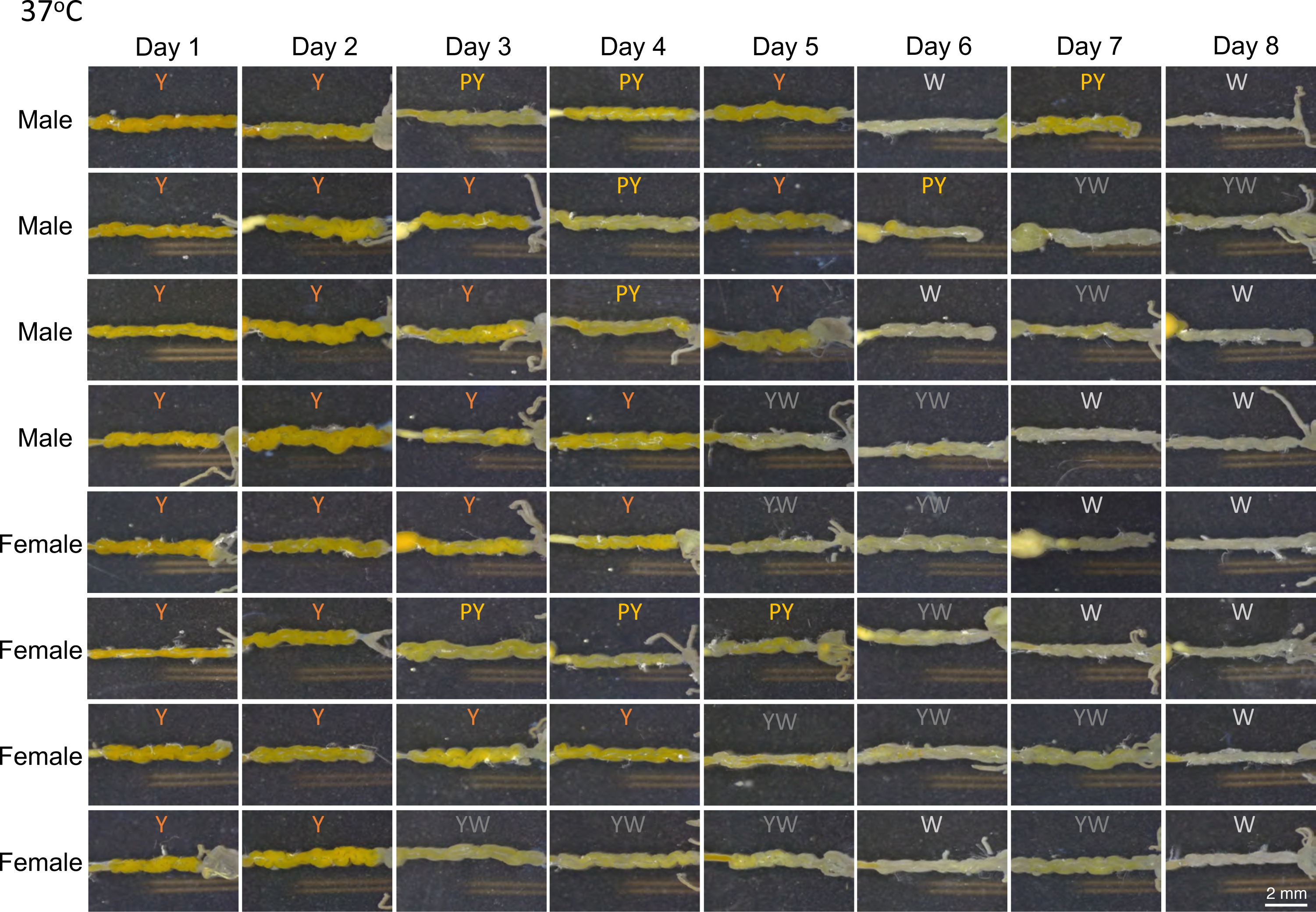
Images of symbiotic organs dissected from *P. stali* adults maintained at 37°C and their color categories abbreviated as Y (yellow), PY (pale yellow), YW (yellowish white) or W (white). For 8 days, 4 males and 4 females were dissected and inspected every day.

**Fig S7.**
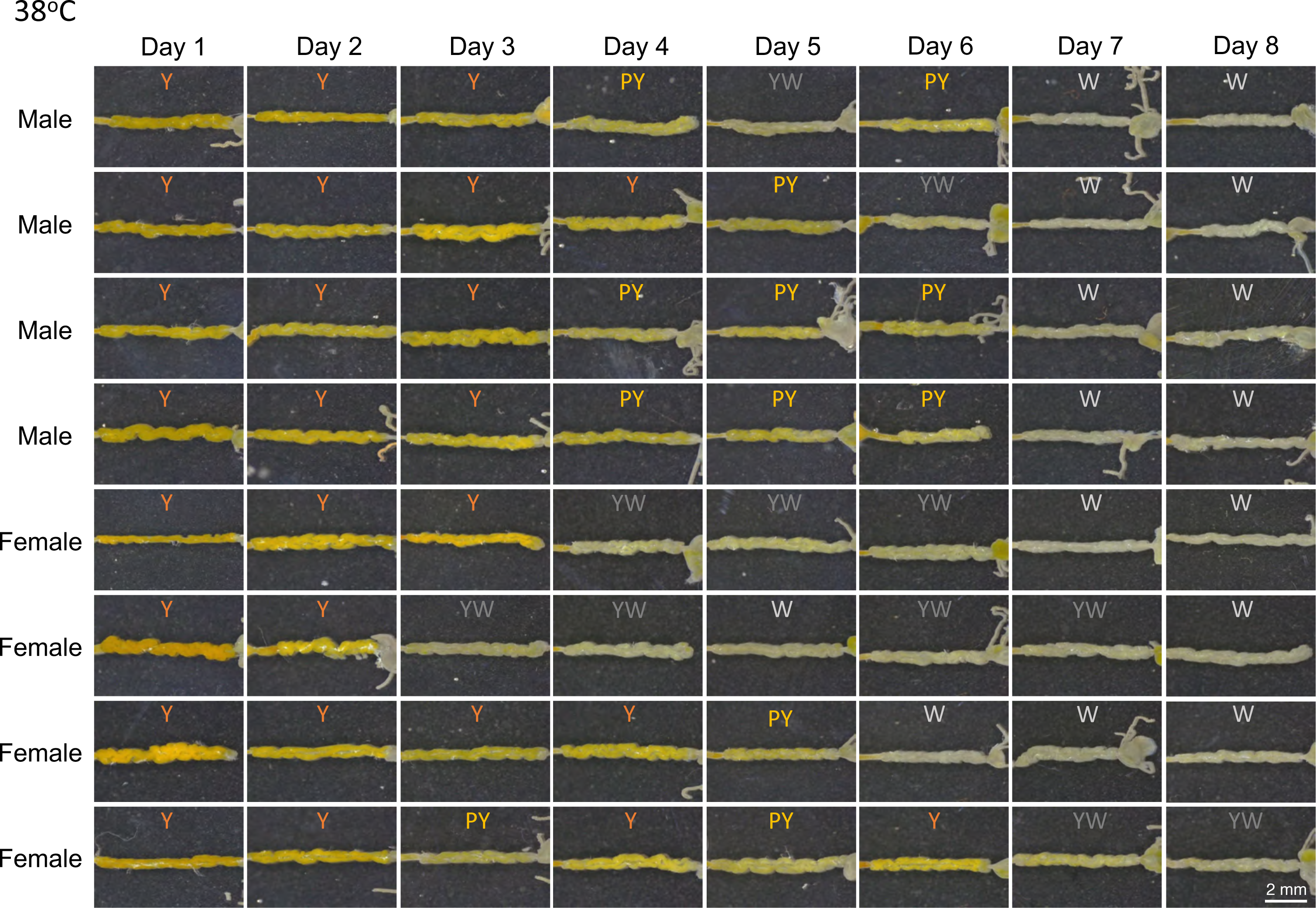
Images of symbiotic organs dissected from *P. stali* adults maintained at 38°C and their color categories abbreviated as Y (yellow), PY (pale yellow), YW (yellowish white) or W (white). For 8 days, 4 males and 4 females were dissected and inspected every day.

**Fig S8.**
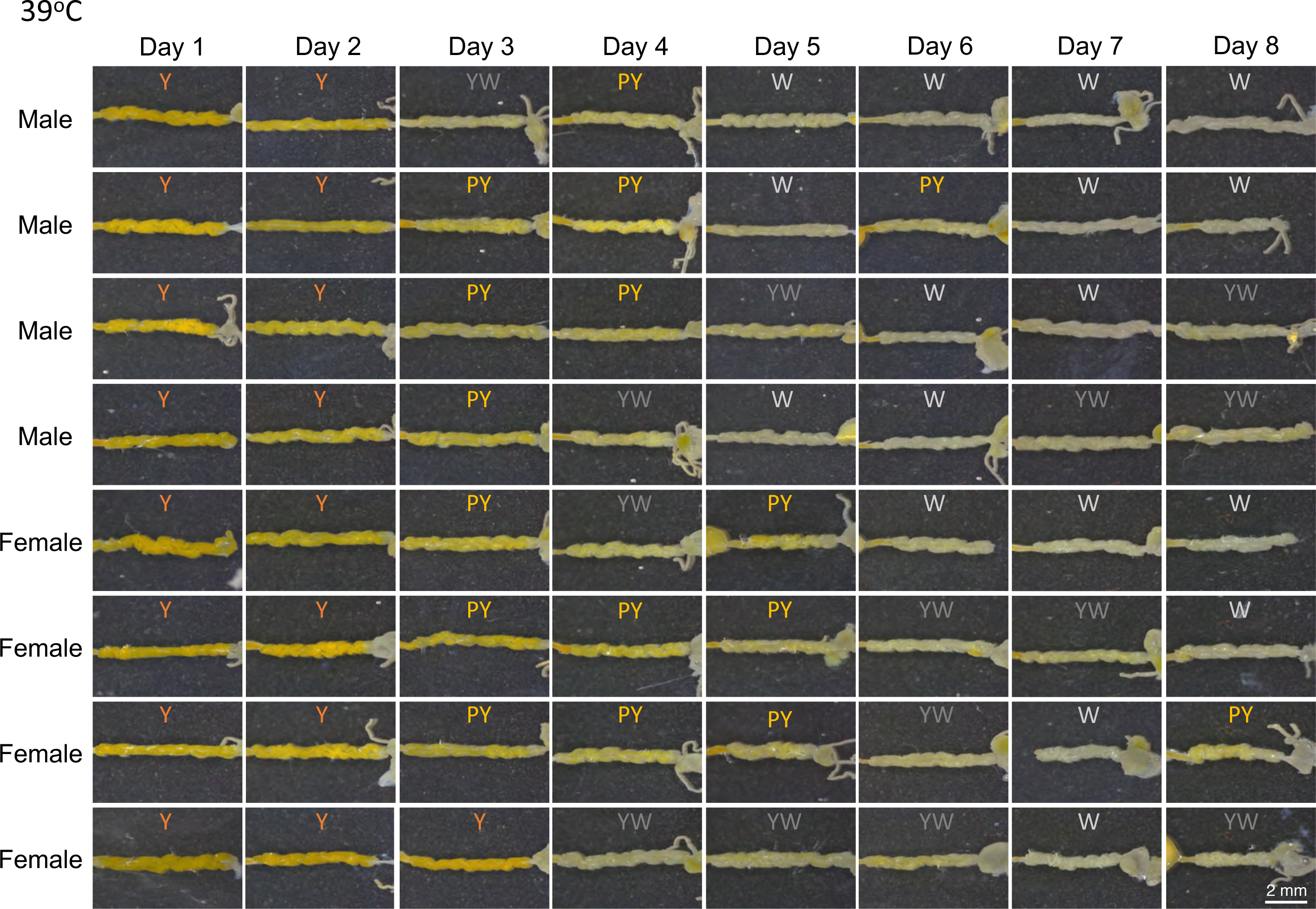
Images of symbiotic organs dissected from *P. stali* adults maintained at 39°C and their color categories abbreviated as Y (yellow), PY (pale yellow), YW (yellowish white) or W (white). For 8 days, 4 males and 4 females were dissected and inspected every day.

**Fig S9.**
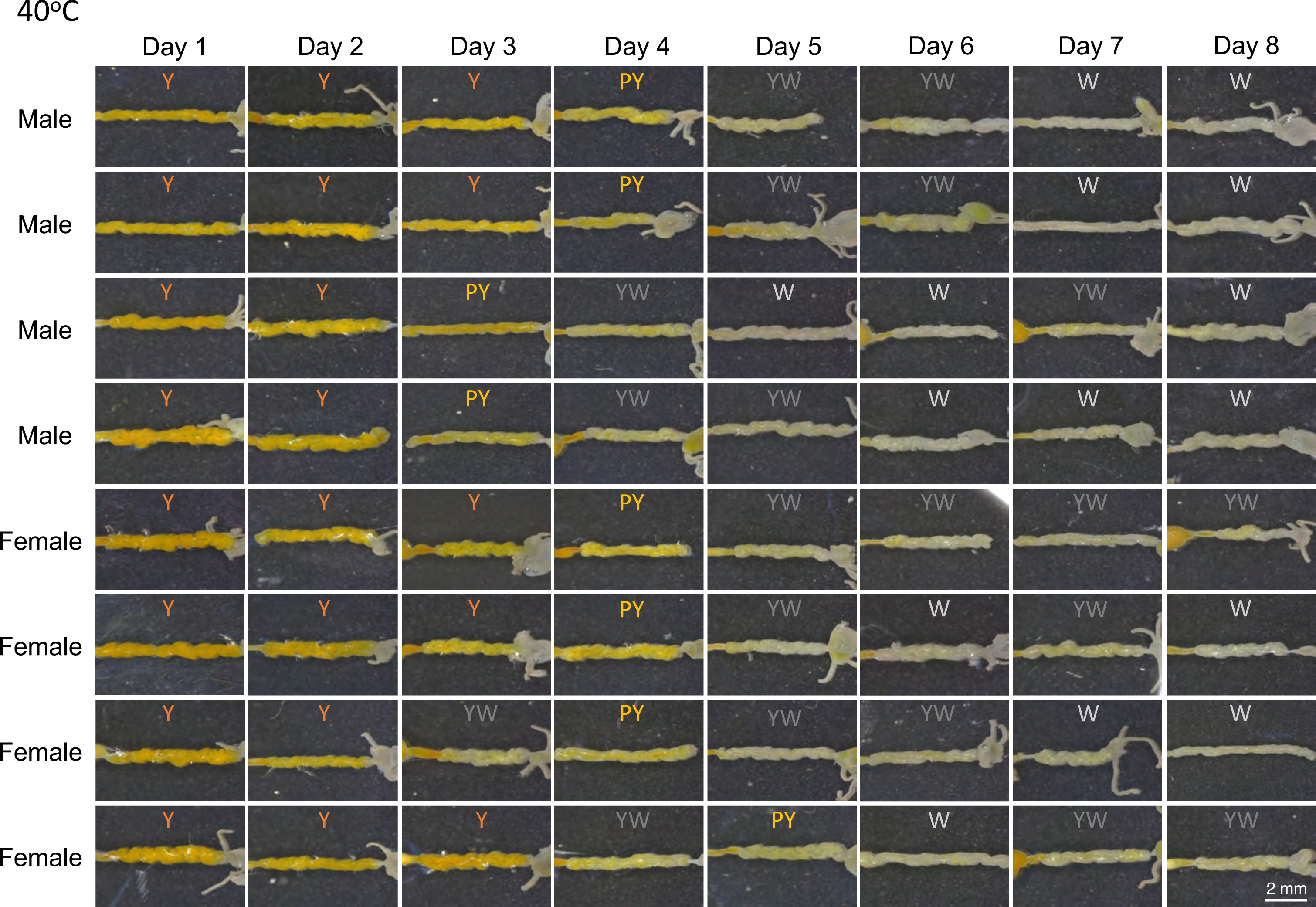
Images of symbiotic organs dissected from *P. stali* adults maintained at 40°C and their color categories abbreviated as Y (yellow), PY (pale yellow), YW (yellowish white) or W (white). For 8 days, 4 males and 4 females were dissected and inspected every day.

